# Riding into the future on a fly: toward a better understanding of phoresy and avian lice evolution (Phthiraptera) by screening bird carcasses for phoretic lice on hippoboscid flies (Diptera)

**DOI:** 10.1101/2021.11.02.466376

**Authors:** Leshon Lee, David J.X. Tan, Jozef Oboňa, Daniel R. Gustafsson, Ang Yuchen, Rudolf Meier

## Abstract

Many phoretic relationships between insects are understudied because of taxonomic impediments. We here illustrate for avian lice riding on hippoboscid flies how new natural history data on phoretic relationships can be acquired quickly with NGS barcoding. Most avian lice are host-specific, but some can arrive on new hosts by riding hippoboscid flies that feed on bird blood. Our summary of the literature yielded 254 published records which we here show to belong to two large and 13 small interaction networks for birds, flies, and lice. In order to generate new records, we then developed a new protocol based on screening bird carcasses sourced by citizen scientists. The inspection of 131 carcasses from Singapore lead to the first record of a *Guimaraesiella* louse species riding on *Ornithoica momiyamai* flies collected from a pitta carcass. Phoresy may explain why this louse species is now known from three phylogenetically disparate hosts (*Pitta moluccensis*, *Ficedula zanthopygia*; *Pardaliparus elegans*). A second new case of phoresy enhances a large interaction network dominated by *Ornithophila metallica*, a cosmopolitan and polyphagous hippoboscid fly species. Overall, we argue that many other two- and three-way phoretic relationships between arthropods (e.g., mites, pseudoscorpiones, beetles, flies) can be resolved using cost-effective large-scale NGS barcoding, which can be used to pre-sort specimens for taxonomic revision and thus partially overcome some taxonomic impediments.

## Introduction

Phoretic relationships between phylogenetically disparate species are common in insects. Small arthropod phoronts regularly use larger arthropod and vertebrate species for transportation. Phoresy can be an important precursor for parasitism/symbioses (Bartlow & Agosta, 2021) and often involve species that deliver important ecosystem services (e.g., decomposition of dung and carrion). Yet, comparatively little is known about the ecology and evolution of phoresy although such relationships are common. Arguably, some of the biggest obstacles for understanding phoresy is identifying which species uses which other species for a ride; i.e., taxonomic impediments. They are often serious because resolving one phoretic relationship (1) requires taxonomic expertise for at least two groups (host and phoront), (2) the phoront tends to be small and thus has a high chance to belong to taxonomically poorly known clade (e.g., mites, lice), and (3) the phoront is often discovered sitting on a host that is studied by a biologist who is mostly interested in the host. This means that most casual observations of phoretic arthropods are either never published or hidden within the literature on the host.

A particularly fascinating, but also slightly atypical phoretic relationship involves three parties: some species of avian lice (Phthiraptera) use blood-sucking flies (Hippoboscidae: Diptera) to travel between avian hosts. All avian lice are flightless obligate ectoparasites that live on birds throughout their life. Louse transmission between hosts usually requires direct physical contact such as parent-offspring interaction (e.g. Clayton & Tompkins, 1994; Brooke, 2010). However, some lice can arrive at new hosts via phoresy on flying insects (Keirans, 1975a) or brood parasitism (Hahn et al., 2000, but see Balakrishnan & Sorenson, 2007). The most important agent are hippoboscid “louse” flies (Diptera: Hippoboscidae), which are highly mobile hematophages. Some avian lice are capable of traveling on these flies by attaching themselves to legs or abdomina (Figure 3A - C). Hippoboscidae is a moderately-sized family of ca. 200 described species of which >80% are bird parasites (Petersen et al., 2007; Dick, 2006) belonging to two clades. Most species little evidence for host specificity and many species have very wide geographic distributions (Bequaert, 1953); i.e., these hippoboscid species have the potential to transfer lice between different bird orders. Add the fact that many birds are migratory and it becomes clear that phoretic lice may be able to jump between distantly related bird species and continents. In comparison, there are only a few records of phoretic interactions between avian lice and other insects such as butterflies and bees (Keirans, 1975a).

Not all avian louse species are involved in phoresy. Phoresy is mostly found among chewing lice (Ischnocera), which feed on feather and skin detritus, while there is only one known case of phoresy by blood-feeding lice in the wild [Amblycera: phoresy of *Hohorstiella giganteus* (Denny, 1842) on an unidentified hippoboscid fly (Hopkins, 1946)]. Even within Ischnocera, phoresy is concentrated in certain ecotypes. For example, ischnocerans specializing on wing feathers of pigeons are better at phoretic attachment than those feeding on body feathers because the latter are more likely to fall off when attempting to ride on a moving fly (Harbison et al., 2008, 2009; Bartlow et al., 2016). This may explain higher levels of genetic structure among pigeon body (non-phoretic) versus wing lice (DiBlasi et al., 2018). Additionally, cophylogenetic analysis comparisons between wing lice and pigeon host versus body lice and pigeon host show that wing lice had higher levels of host switching compared to body lice which generally coevolved with their hosts (Clayton and Johnson, 2003). Body lice were also found to have higher genetic divergence than wing lice, possibly due to the availability of (or rather lack of) hippoboscid flies for phoretic dispersal (Sweet and Johnson, 2018).

However, despite its potential biological importance and recent experimental work (Harbison et al., 2008, 2009; Bartlow et al., 2016), phoresy remains poorly documented in the wild, so that its importance is hard to assess.

We here summarize the literature on louse phoresy and construct species interaction networks. This revealed three major problems. Firstly, the number of observations that provide precise taxonomic information for all species (i.e., bird, fly, and louse) was very limited because this requires extensive taxonomic knowledge for three very different animal groups. Secondly, the number of new records is in steep decline with the majority of published records concentrated in the first half of the 20^th^ century (Keirans, 1975b; Bartlow et al., 2016). This is correlated with the overall decline in the number of natural history publications (Tewksbury et al., 2014). Lastly, a large proportion of the known phoresy records come from temperate Europe and North America despite the comparatively small Holarctic avian fauna (Figure 1).

**Figure 1:**
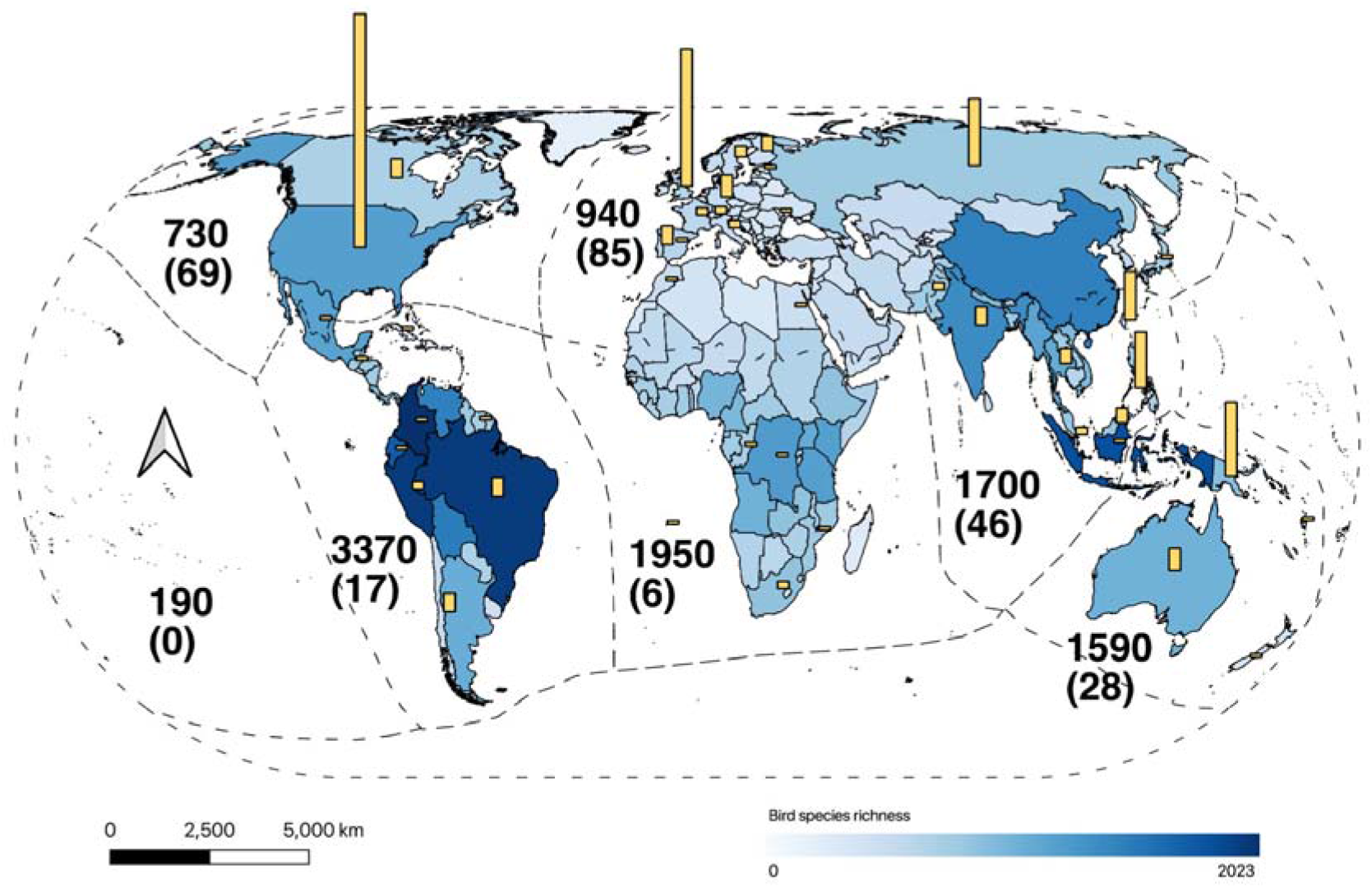
Map of bird species richness (blue shading across countries) and phoresy records (yellow bars) by country illustrating the Holarctic bias of bird-louse-hippoboscid phoresy records. Numbers indicate approximate bird species richness of each biogeographical realm and number of phoresy records (in parentheses). Bird species richness map: Clements et al. (2019), Species richness figures: Newton & Dale 2001, basemap from thematicmapping.org, biogeographical realm shapefile from UNEP-WCMC (2011), generated by QGIS v2.18 (Las Palmas).

We then address these problems by developing a new protocol for obtaining new phoresy records. It is here developed for avian lice, but the molecular techniques are equally applicable for resolving other phoretic relationships. In our study of avian lice, we screened bird carcasses that were sourced by encouraging members of the public to report dead birds in Singapore. This yielded 131 bird carcasses (54 species) from which lice, hippoboscid flies, and mites were collected. For the lice and flies, we address partially the taxonomic impediments that interfere with so much natural history research via NGS barcoding (Wong et al., 2017; Yeo et al., 2018; Srivathsan et al., 2019, 2021). The specimens were sorted to putative species based on these barcodes which can now be obtained at low cost for thousands of specimens within days (Srivathsan et al., 2018, 2019, 2021; Wang et al., 2018; Yeo et al., 2018). Based on this protocol, we here report two new cases of phoresy which add to the species interaction networks identified based on literature data.

## Materials & Methods

### Literature review

Using the records in Bartlow et al. (2016) as starting point, we checked the literature on louse-hippoboscid phoresy for overlooked or misattributed records. In addition, we conducted a literature search using the keywords “Hippoboscid*”, “Mallophaga*”, “Phthiraptera*”, “Louse”, “Lice”, and “Phore*” in Web of Science, Biodiversity Heritage Library, Phthiraptera.info database, and Google Scholar.

### Collection of avian ectoparasites

We collected bird carcasses reported by citizen scientists in Singapore as part of a long-term project to monitor avian mortality due to window-collisions or road accidents (Low et al., 2017; Tan et al., 2017). Carcasses collected between 2013 and 2019 were bagged separately and stored at −20°C. We brushed 131 bird specimens (54 species) for lice and hippoboscid flies using a toothbrush and preserved all ectoparasites in 95% ethanol at −20°C. We then identified hippoboscid flies carrying lice using a Nikon SMZ460 stereomicroscope.

### Ectoparasite DNA barcoding

For all the lice and hippoboscids collected, we determined the number of species-level units using NGS barcodes (Baloğlu et al., 2018; Meier et al., 2016; Wang et al., 2018; Yeo et al., 2020). We extracted genomic DNA from hippoboscid flies and lice using a modified hotSHOT protocol (Truett et al., 2000). For lice, we used 10μl of alkaline lysis buffer and neutralizing reagent per specimen, while the quantities were increased to 15μl for hippoboscids. A 313-bp Cytochrome Oxidase 1(COI) minibarcode was amplified using modified primers published in Geller et al. (2013) and Leray et al. (2013) [m1COlintF: 5’-GGWACWGGWTGAACWGTWTAYCCYCC-3’ (Leray et al., 2013) and modified jgHCO2198: 5’-TANACYTCNGGRTGNCCRAARAAYCA-3’ (Geller et al., 2013)]. In addition, we amplified another 379-bp COI fragment for one specimen for all putative louse species to ensure overlap with the barcoding region used in previous louse barcoding studies (primers: L6625F: 5’-CCGGATCCTTYTGRTTYTTYGGNCAYCC-3’ and H7005R: 5’-CCGGATCCACNACRTARTANGTRTCRTG-3’: Hafner et al., 1994). For all amplicons, the forward and reverse primers were tagged with a 9-bp oligonucleotide tag at the 5’-end. Unique tag combinations could then be used to distinguish the amplicons for each specimen (Meier et al., 2016; Wang et al., 2018). Successful amplification was checked for a subsample of all PCR reactions on a 1.5% agarose gel. Subsequently, all PCR products were pooled and purified using Sera-mag SpeedBeads (Fisher Scientific) as per Rohland & Reich (2012). The pooled and cleaned amplicons used in our report on phoresy were sequenced at the Genome Institute Singapore on a partial Illumina Hiseq2500 Rapid Run lane (251-bp paired-end).

We used default parameters in PEAR 0.9.6 (Zhang et al., 2014) to merge paired-end reads and demultiplex the reads via a Python script utilizing the unique F and R primer tag combinations for each specimen (Meier et al., 2016; Wang et al., 2018). Read counts and variants were processed according to the quality control pipeline described in Meier et al. (2016). The read variants with the highest and second-highest counts for each specimen were checked against GenBank using BLAST and reads with similarity scores >97% to non-Phthiraptera taxa were removed. We then aligned all 313bp COI barcode sequences using Mafft v7 (Katoh & Standley, 2013) and clustered the barcodes by pairwise genetic distances using a 2%, 3%, 4% and 5% distance thresholds (Meier & Wheeler, 2008; Meier et al., 2008). This allowed for identifying stable Molecular Operational Taxonomic Units (MOTUs).

### Identification of Specimens

We used BLAST to obtain species-level identifications for the louse and fly barcodes (>97%). In addition, we used morphology to confirm species limits and to identify the specimens based on keys and checklists (Maa, 1966a; 1966b; 1969a; 1969c; 1969d; Gustafsson & Bush, 2017). We documented phoresy and the morphology of the ectoparasites by obtaining high-resolution dorsal and ventral views imaged at different focal lengths with a Dun Inc. Passport II Imaging system (Canon 7D Mk II with MPE-65 lens). Images were then focus-stacked using Zerene Stacker (Zerene Systems LLC) and prepared for publication using Adobe Photoshop CS5.

## Results

### Literature Review

We found 254 literature records (1857–2021) of louse-hippoboscid phoresy with at least a genus-level identification for either lice or hippoboscids (Table S1). Three records reported in Bartlow et al. (2016) were misattributions and two records could not be verified based on primary sources (see Table S1). All louse-hippoboscid phoresy records are shown in Figure 2. The species interaction network omits records pertaining to louse-bird interactions only except for the newly discovered louse-bird interactions from Genbank and omits records that do not have species level identification for the louse, fly, or bird except for the louse species involved in the newly discovered phoresy interaction.

**Figure 2:**
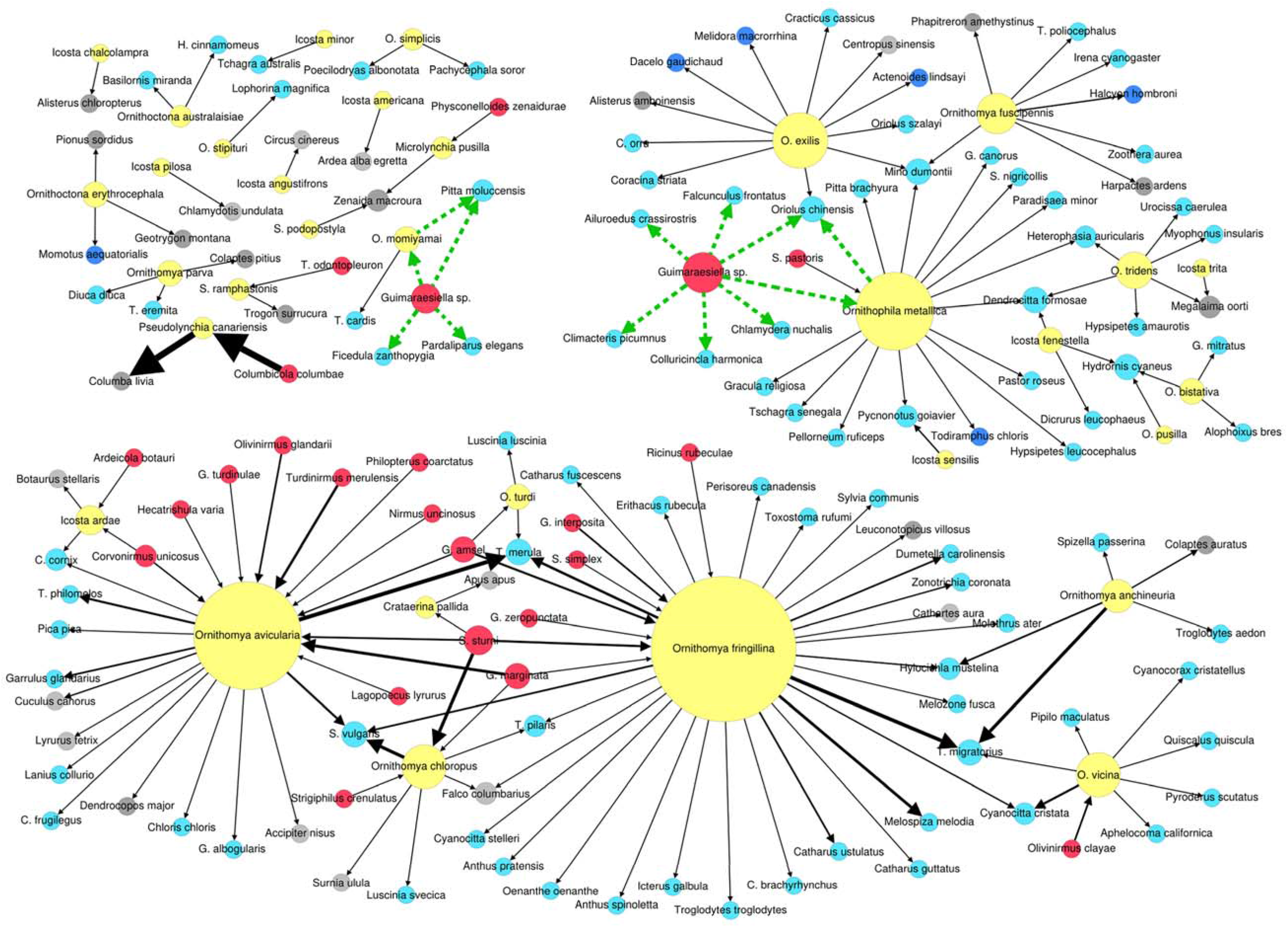
Species interaction network of phoretic avian lice records (red) on hippoboscid flies (yellow), and associated birds (light blue=Passeriformes; dark blue=Coraciiformes; grey=remaining bird orders). Size of nodes corresponds to interaction number. Genus abbreviations for birds: C.=*Corvus*, G.=*Garrulax*, H.= *Hypocryptadius*, S.=*Sturnus*, T.=*Turdus*; for flies: O.=*Ornithoica*, S.=*Stilbometopa*; lice: G.=*Guimaraesiella*, S.=*Sturnidoecus*, T.=*Trogoninirmus*). Thickness of black arrows corresponds to the number of records for the particular interaction. Green dashed arrows refer to the newly discovered interactions including louse-bird interactions from Genbank.

### New louse-hippoboscid phoresy records

We collected 32 hippoboscid flies from 22 of the 131 bird carcasses (13 bird species). Of the 32, three carried phoretic lice (Figure 2 and Table S1). Two hippoboscid specimens (ZRC_BDP0273056, ZRC_BDP0273057) collected from a carcass of a Blue-winged pitta (*Pitta moluccensis* (Müller, 1776) specimen code CR465), carried three phoretic lice in total (Louse specimens ZRC_BDP0298043 and ZRC_BDP0298044 attached to hippoboscid specimen ZRC_BDP0273056. Louse specimen ZRC_BDP0298045 attached to hippoboscid specimen ZRC_BDP0273057). The third hippoboscid specimen (ZRC_BDP0273050) carried four phoretic lice (ZRC_BDP0298039, ZRC_BDP0298040, ZRC_BDP0298041, ZRC_BDP0298042) and was obtained from a Black-naped oriole (CR619: *Oriolus chinensis* Linnaeus, 1766). We also barcoded the free-roaming lice on the bodies of the Blue-winged pitta and the Black-naped oriole to determine whether the lice belonged to the phoretic species.

### NGS barcoding of lice and hippoboscids

603 louse specimens (including the phoretic lice) and all 32 hippoboscid specimens were successfully barcoded (Genbank Accession numbers for specimens involved in phoresy: MT762409-MT762417). Clustering the 603 louse barcodes using Objective Clustering showed that the number of louse MOTUs was stable at 56 MOTUs for pairwise distance (p-distance) thresholds between 2–5%. For hippoboscids, the number of MOTUs obtained using Objective Clustering was 12 at 2–3%, and 11 between 4–5%. 2–5% p-distance thresholds were based on literature (Meier et al., 2006). Using Assemble Species by Automatic Partitioning (ASAP: p-distances model) (Puillandre et al., 2021), the top partition clustered the louse and hippoboscid sequences into 57 (ASAP score = 4) and 12 MOTUs (ASAP score = 1.5) respectively. The phoretic lice belonged to two louse species, which clustered with other free-roaming lice from the same bird carcass. The phoretic louse species from the Blue-winged pitta is also known from a Yellow-rumped flycatcher carcass [*Ficedula zanthopygia* (Hay, 1845); Table S1] when compared to other data from Singapore (Lee, unpublished data). However, the same louse species was not detected on two additional Blue-winged pitta carcasses with lice (Lee, unpublished data). The second phoretic louse species reported here was found on one Black-naped oriole carcass.

### Identification of flies and lice

None of the 313bp COI barcodes for lice had species-level matches, but we obtained a 99.74% match for the 379-bp sequence obtained from a Blue-winged pitta louse for *Brueelia* sensu lato in Genbank, although the species is now assigned to the genus *Guimaraesiella* (Gustafsson & Bush, 2017). The sequence was obtained from an Elegant Tit (*Pardaliparus elegans* Lesson, 1831) in the Philippines (Johnson et al., 2002b; Bush et al., 2016; GenBank Accession number: AY149382). The Black-naped Oriole louse had five Genbank matches >97% to a 379-bp sequence of a different louse species identified as *Brueelia* sensu lato. This species has now also been re-assigned to the genus *Guimaraesiella*. One match (98.17%) was obtained from a louse on a Brown treecreeper (*Climacteris picumnus* Temminck & Laugier, 1824), another (97.38%) from a louse on a Green catbird [*Ailuroedus crassirostris* (Paykull, 1815)], and three matches (97.13%) from lice on a Crested shriketit [*Falcunculus frontatus* (Latham, 1801)], a Grey shrikethrush [*Colluricincla harmonica* (Latham, 1801)], and a Great bowerbird (*Chlamydera nuchalis* Jardine & Selby, 1830). Of the flies, only five 313bp COI barcodes had species-level (>97%) matches to *Pseudolynchia canariensis* (Macquart, 1839). The hippoboscids on the Blue-winged pitta were thus identified using morphological keys as *Ornithoica momiyamai* Kishida, 1932 (Figure 3A & 3B) and the one on the Black-naped oriole as *Ornithophila metallica* (Schiner, 1864) (Figure 3C).

**Figure 3:**
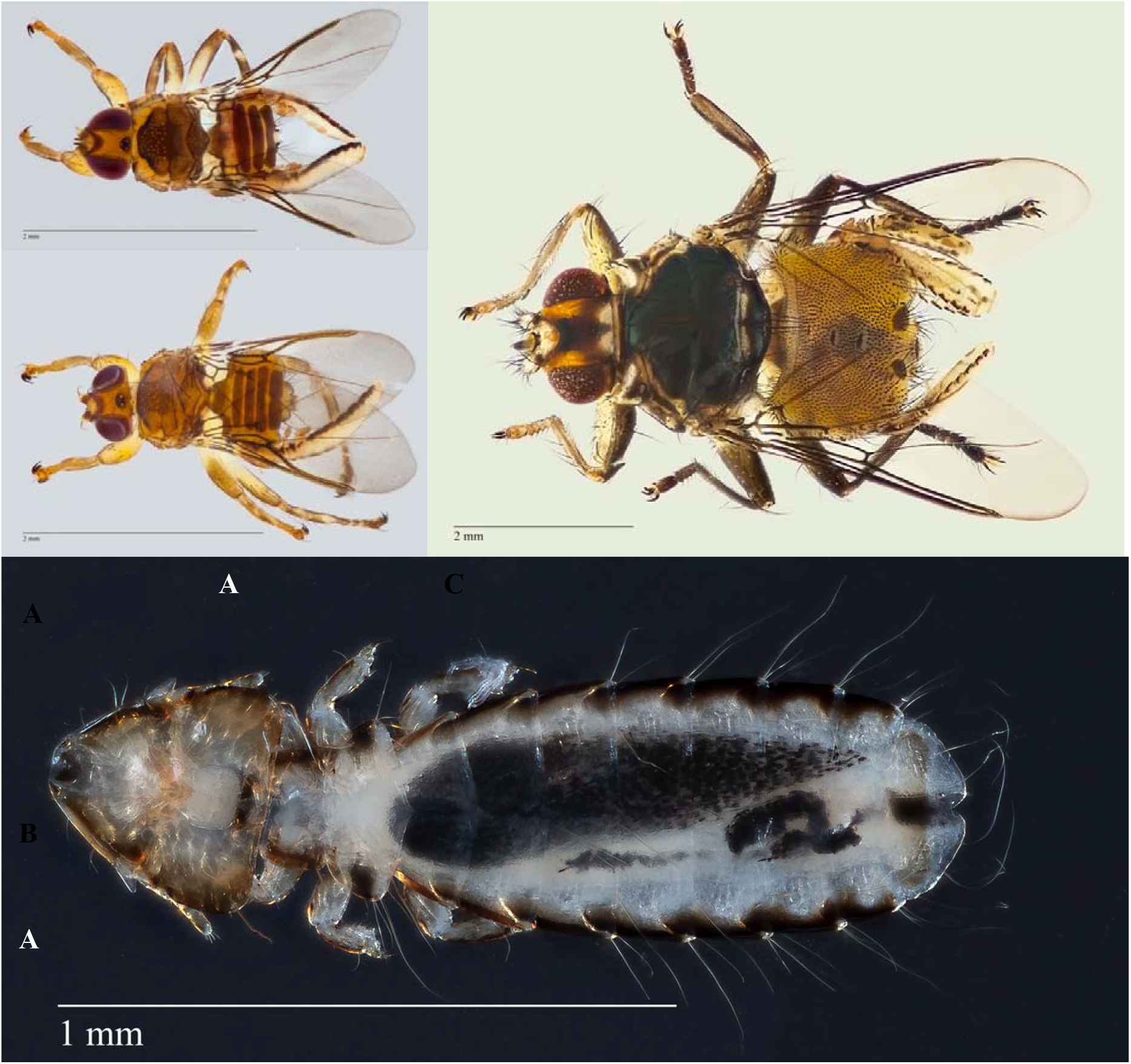
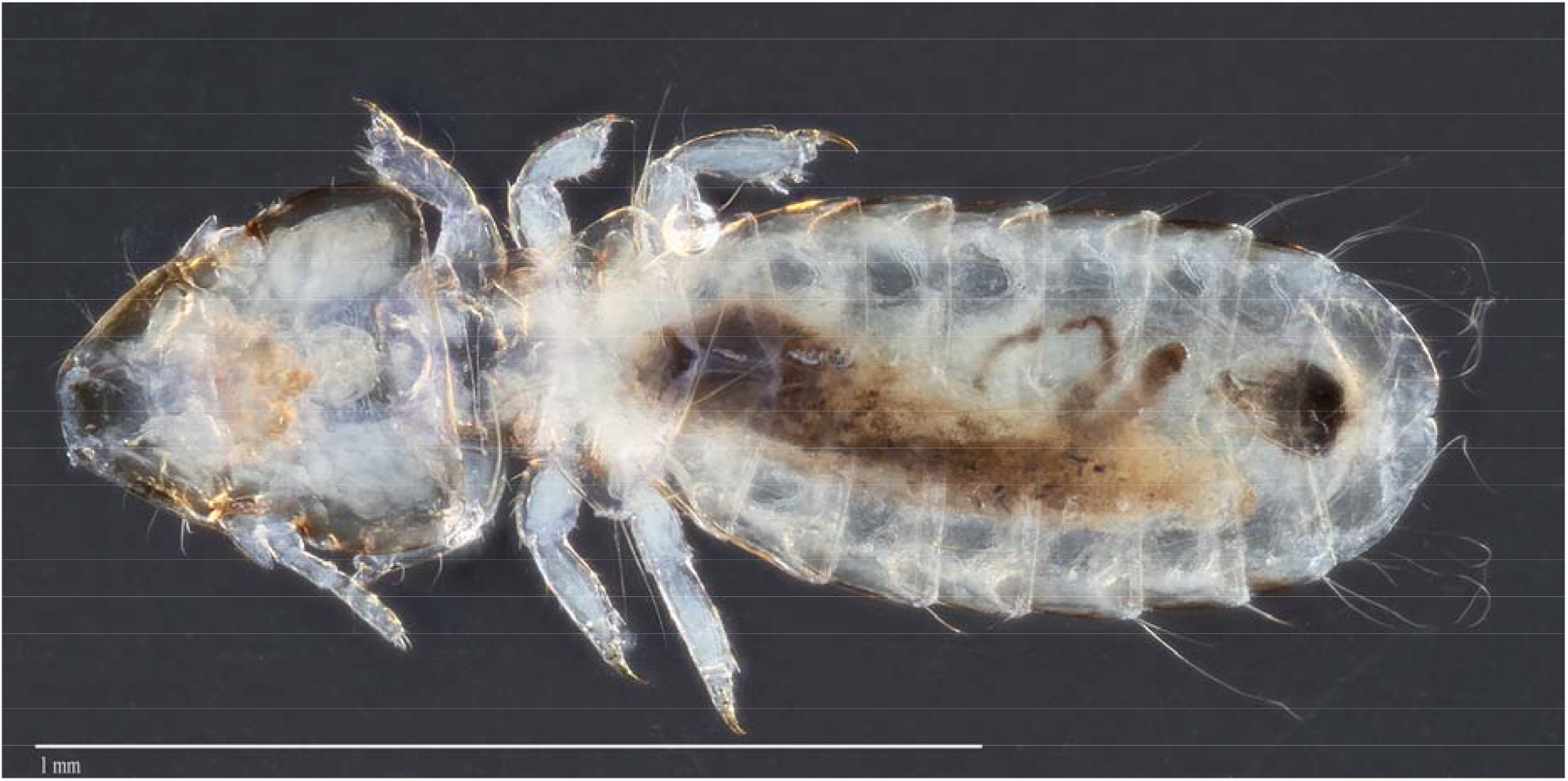
(A-B): *Ornithoica momiyamai* (ZRC_BDP0273056) with *Guimaraesiella* specimens attached to abdomina. (C): *Ornithophila metallica* (ZRC_BDP0273050) with four *Guimaraesiella* specimens attached to its abdomen. (D): Habitus of *Guimaraesiella* ZRC_BDP0298043. (E): Habitus of *Guimaraesiella* ZRC_BDP0298039.

### New Cases of Phoresy

Based on morphology we identified the phoretic lice found on the flies (*Ornithoica momiyamai*) obtained from the *Pitta moluccensis* carcass as belonging to the “core” group of the genus *Guimaraesiella* ZRC_BDP0298043 (*sensu* Gustafsson et al., 2019a). The genus was previously not known to feed on pittas (Figure 3D; Somadder & Tandan, 1977; Gustafsson & Bush, 2017). It is now known to be shared between Blue-winged pittas, Yellow-rumped flycatchers, and the Philippine endemic Elegant Tit. The louse specimen from the Elegant Tit is genetically very similar to lice from at least 24 other host species (Bush et al, 2016), and morphologically indistinguishable from specimens from additional species (D.R. Gustafsson, unpublished data). Based on morphology, we identify the phoretic louse on the fly *Ornithophila metallica* from the *Oriolus chinensis* as also belonging to the genus *Guimaraesiella* ZRC_BDP0298039 (Figure 3E), which matched on Genbank with lice recovered from five Australasian passerine species.

## Discussion

Phoresy remains an understudied and underappreciated phenomenon because taxonomic impediments are particularly likely to affect the chance that new observations are communicated. We here propose to address the taxonomic impediments by sorting specimens to species-level based on NGS barcodes. We believe that NGS barcodes are a good choice (Srivathsan et al., 2018, 2019; Wang et al., 2018; Yeo et al., 2018), because they are now cost-effective (<US$0.10 per specimen: Srivathsan et al., 2021; Yeo et al., 2021) and can be obtained with a very basic laboratory setup within days (Srivathsan et al., 2018, 2019; 2021; Wang et al., 2018). Barcodes can be easily matched across samples collected at different times and places so that host and phoront species repeatedly involved in a phoretic relationship can be prioritized for identification and/or description. In addition, specimens, especially in their early instar stages, can be assigned to putative species via barcode databases (Yeo et al., 2018). We here used NGS barcodes to find new three-way species interactions between louse, fly, and bird, but the same techniques are also very valuable for ornithologists who are interested in two-way relationships between birds and their flies or lice. The same techniques are also very valuable to address numerous other phoretic relationships which often involve one partner that belongs to a clade that is taxonomically particularly poorly known.

Some biologists may object that using barcodes for delimiting putative species is unsatisfactory, but we would argue that it is a step in the right direction as documented by the scarcity of species-level louse identifications in the published literature. They were only available for less than a third of the published cases and an additional ~20% of lice were only identified to genus. If one wanted to improve the taxonomic resolution of these records, the relevant specimens would have to be located in numerous collections. We predict that many would not be found and/or not sufficiently well preserved for identification based on morphology. Such species-level matching is more straightforward with barcodes. For example, matching the *Guimaraesiella* record from Singapore to a louse record from the Philippines obtained almost 20 years ago took minutes via NCBI although the genus attribution of both the louse and host species in question had changed (Del Hoyo et al., 1992; Gustafsson & Bush 2017). Note, however, that such assignments of specimens to putative species via barcodes is no endorsement for describing species based on barcodes only. As argued by numerous authors (Ahrens et al., 2021; Engel et al., 2021; Meier et al., 2021), barcode clusters are first-pass grouping statements that require additional testing before they can be properly described as species. Instead, pre-sorting specimens to putative species using barcodes is the first step toward taxonomic revision. NGS barcoding is now fast and cost-effective enough that it can replace morphological pre-sorting. If all specimens are barcoded, it also yields approximate abundance information and morphological testing of species boundaries can be carried out at the haplotype level (Hartop et al. 2021).

The impact of taxonomic impediments on understanding phoresy is illustrated by our literature review. Many records could not be used because they lacked taxonomic resolution – mostly for lice. This meant that a comparatively low number of reports cover louse-hippoboscid phoresy in sufficient detail to assess the importance of phoresy for louse-host specificity and the co-speciation between avian lice and bird hosts. This is unfortunate given that bird lice are species-rich (Price et al., 2003; Gustafsson et al., 2019b) with many bird species hosting several louse species which makes it likely that the number of louse species will eventually exceed the number of bird species. Despite the small number of published records, we were able to reconstruct two large species interaction networks that connect 87 species of birds, 16 species of flies, and 18 species of lice. The networks are so sizable because many hippoboscid flies visit a very large number of hosts and have wide geographic distributions (Bequaert, 1953). In addition, many bird hosts are migratory which increases the chance of transferring lice from birds breeding in the temperate region to birds resident in the tropics and vice versa. Note that the networks would be even larger if we had mapped all two-way species interactions (bird-louse and fly-bird). For example, the *Guimaraesiella* specimen obtained from the Blue-winged Pitta belongs to an undescribed species that has been found on 24 other host species (see Bush et al., 2016). Based on morphology, specimens likely belonging to the same species are furthermore known from another 30+ host species (D.R. Gustafsson, unpublished data). The geographic range of all these specimens spans from New Guinea and Australia over China, Thailand and India to Malawi (Bush et al., 2016). This illustrates that screening a sufficiently large number of bird carcasses and barcoding the phoronts would likely yield even more impressive and large species interaction networks.

How species interaction networks expand with the addition of only a few new records is illustrated by the new phoresy cases reported here. One involves two hippoboscid flies belonging to the same species (*Ornithoica momiyamai*) obtained from the same *Pitta moluccensis* carcass. Both carried the same louse species (*Guimaraesiella* sp.) that we had previously already found on a Yellow-rumped Flycatcher carcass from Singapore. As a long-range migrant, Yellow-rumped Flycatchers breed in East Asia and overwinters in western Sundaland (Del Hoyo et al., 1992). This makes it likely that it is visited by a large number of hippoboscid flies throughout its range which may result in the transmission of lice belonging to the *Guimaraesiella* species to numerous bird species (see Bush et al., 2016). It may also explain why this *Guimaraesiella* species was previously genotyped from a Philippine endemic, the Elegant Tit (Del Hoyo et al., 1992). Finding the putatively same *Guimaraesiella* species on a pitta in Singapore was unexpected because it may represent a case of an incipient phoresy-mediated host-switch given that no other *Guimaraesiella* species has ever been reported from a pitta species (Gustafsson & Bush, 2017) and we did not find *Guimaraesiella* specimens on two additional pitta carcasses with lice. Pittas are typically parasitised by lice of the genus *Picicola* (Somadder & Tandan, 1977) which are only distantly related to the *Brueelia*-complex, of which *Guimaraesiella* is part. Of course, additional records would be welcomed, given that it is conceivable that the *Guimaraesiella* species may not be able to establish itself on the pitta.

The *Guimaraesiella* species was found on a hippoboscid species (*Ornithoica momiyamai*) that is found throughout Asia and is known to feed on 11 bird species (Maa, 1969; Suh et al., 2012). This is a small number of bird species compared to the hippoboscid species involved in the second new case of phoresy reported here. It involves a *Guimaraesiella* louse species clinging to an *Ornithophila metallica* fly. This fly species has an exceptionally wide distribution across all biogeographic regions (except Antarctica) and is known to feed on bird species belonging to 134 genera (Maa, 1969; Suh et al., 2012). It is thus surprising that the *Guimaraesiella* louse species that uses *Ornithophila metallica* does not have a wider distribution. Our louse specimen is most similar to specimens from Australasia (see Bush et al., 2016) and our record is the first outside of this region although riding on *Ornithophila metallica* should open up many additional opportunities for host switching. Within Australasia, this louse species is known from non-migrating hosts from at least four different families, suggesting that phoresy may have been important for the host distribution.

We obtained the new records by screening >100 bird carcasses sourced with the help of citizen scientists. We found that ~73% of the carcasses had lice, ~16% had hippoboscid flies, and ~1.5% had hippoboscid flies carrying lice. Louse phoresy on hippoboscid flies is thus not particularly common, but may nevertheless be a significant phenomenon over evolutionary time given that recent studies of chewing lice have shown that successful establishment of louse populations on distantly related hosts may be rare, but not impossible (Sychra et al., 2014; Bush et al., 2016; Gustafsson & Bush, 2017; Gustafsson et al., 2019c). Obtaining the new phoresy records was not time consuming because finding hippoboscid flies on bird carcasses is fast and the number of birds that are killed annually is vast: Loss et al. (2014) estimate that between 365 and 988 million birds die from window collisions in the United States alone and Grilo et al. (2020) estimate 194 million annual bird road kills in Europe. In Singapore, Tan et al. (2017) collected 104 non-migratory and 204 migratory bird carcasses between 2013 to 2017 despite avoiding common species. We predict that hundreds of new phoresy records could be obtained via carcass screening within a short time period, especially if additional fly specimens were sourced from bird ringing initiatives. Screening bird carcasses for ectoparasites is thus a cost-effective complement to field sampling. It is non-invasive and requires very little specialist knowledge or field expertise. Note, however, that many flies abandon birds once the carcass cools to room temperature (Bequaert, 1953), so that screening has to be quick and thus be made part of carcass salvage protocols.

The widespread distributions and unfastidious taste of many hippoboscid flies explains why two of our species interaction networks are large although most published records lack taxonomic resolution for lice (Table S1 and Figure 2). Taxonomic impediments interfere with the publication of many natural history observations (Srivathsan et al. 2019) and are particularly severe here, because it is difficult to assemble a team consisting of on ornithologist and two entomologists with complementary taxonomic expertise. Even if such a collaboration can be arranged, the lack of comprehensive keys and the large number of undescribed louse species are major obstacles. Yet, obtaining accurate species-level data is crucial for reconstructing species interaction networks and understanding how the relationships between bird hosts, avian lice, and how hippoboscid flies can shape louse evolution.

## Conclusions

Our study identifies significant gaps in our understanding of the phoretic relationships between avian lice, hippoboscids, and birds. The gaps are particularly large for many areas with rich avian faunas. At this point, we do not know whether these gaps reflect biology or sampling bias, but this question could be addressed quickly via screening bird carcasses from different biogeographic regions. This will not only facilitate the study of bird-louse coevolution as mediated by hippoboscid flies, but also help with understanding why certain species/genera of flies and lice are frequently involved in phoretic relationships. It is important to restart natural history research (Tewksbury et al., 2014). Fortunately, combining traditional techniques such as carcass screening with new molecular techniques greatly facilitate and improve our ability to obtain new and species-level results (Wong et al., 2017; Yeo et al., 2018; Srivathsan et al., 2019).

## Acknowledgements

We thank Kim Mi Jin for her help with the translation of Japanese and Korean papers and Tommy Tan, Goh Poh Moi, Morgany D/O Thamgavelu, and Ismail Bin Arshad for providing us access to NUS Lab 7’s equipment and space for ectoparasite collection. The research was supported by a Ministry of Education grant on biodiversity discovery (R-154-000-A22-112).

